# Effect of endogenous and exogenous ketosis in acute spinal cord injury in Wistar rats

**DOI:** 10.64898/2026.02.09.704895

**Authors:** A. Cervantes, P. Montiel-de la Rosa, A. Soto-Mota, J. Alanis-Mendizabal, M. Valderrama, V. Ramírez, C. Bautista, R. Vicuña, G. Reyes-Soto, M.A. Pineda-Castillo, M.G. Palacios-Saldaña, C.C. Bravo-Reyna

**Affiliations:** Departamento de Investigación Experimental y Bioterio. Instituto Nacional de Ciencias Médicas y Nutrición Salvador Zubirán. Mexico City, Mexico; Unidad de investigación en enfermedades metabólicas. Instituto Nacional de Ciencias Médicas y Nutrición Salvador Zubirán. Mexico City, Mexico; Departamento de Nutrición Animal. Instituto Nacional de Ciencias Médicas y Nutrición Salvador Zubirán. Mexico City, Mexico; Departamento de Reproducción. Instituto Nacional de Ciencias Médicas y Nutrición Salvador Zubirán. Mexico City, Mexico; Departamento de Patología, Hospital Central Sur de Alta Especialidad PEMEX, Mexico City, Mexico; Unidad de neurocirugía oncológica. Instituto Nacional de Cancerología, Ciudad de México, Mexico

**Keywords:** Spinal cord injury, Ketosis, Methylprednisolone, Inflammation, Oxidative stress, 1, 3 hydroxybutanediol

## Abstract

Acute spinal cord injury is a condition with a poor prognosis, leading to reduced quality of life and high economic costs for both patients and health systems, and only a minority of patients with this injury can achieve a meaningful recovery. Given the high frequency of traumatic events such as vehicle collisions and falls, research aimed at limiting injury extension, promoting neuronal recovery, and improving prognosis is essential. It has been shown that ketone bodies have anti-inflammatory properties and also mediate the Nrf2 pathway, exerting antioxidant effects. The aim was to identify any alternative to mitigate the extension and progression of secondary injury by using 1,3 butanediol as a β-hydroxybutyrate source. An experimental model (n= 60) of clinically healthy male Wistar rats were used, divided into five groups (n=12) as a control group, while the remaining rats were subjected to extradural spinal cord clipping according to the following groups (n=12): Spinal Cord Injury (SCI); endogenous ketosis + methylprednisolone (Endo-K+MP); exogenous ketosis + methylprednisolone (Exo-K+MP), and methylprednisolone (MP). After 8 hours of spinal cord injury, tissue was collected, immunohistochemistry and PCR analyses were carried out. Nrf2 and 3NT for the antioxidant pathway, and HIF-1, NFkB, NLRP3, TNF-ɑ and IL-1β for inflammation were analyzed. Results demonstrated that the groups thrown into a ketosis state had better outcomes, and, according to the Exo-k+MP and Endo-k+MP groups, increased Nrf2 and decreased 3NT (p < 0.05), which resulted in an upregulation of antioxidant pathways. According to HIF-1α and NFκB, Endo-K+MP showed better outcomes p<0.05 and proinflammatory cytokines showed the same pattern as the standard treatment (MP) p<0.05). Overall, our results also demonstrated a downregulation of inflammatory pathways.

**Author Summary:** Spinal cord injury is a devastating condition that frequently results in permanent neurological damage, reduced quality of life, and high social and healthcare costs. Current treatments are limited and mainly focus on reducing inflammation after injury, with methylprednisolone being one of the most commonly used therapies despite its associated adverse effects. Previous studies have shown that ketosis, a metabolic state characterized by increased ketone bodies, has anti-inflammatory and antioxidant properties in several neurological conditions.

In this study, we investigated whether inducing ketosis could improve early outcomes after acute spinal cord injury. Using a rat model, we compared the effects of endogenous ketosis (induced by fasting) and exogenous ketosis (induced by ketone precursors), alone or in combination with methylprednisolone. We analyzed inflammatory and oxidative stress pathways using immunohistochemistry and molecular techniques.

Our findings show that ketosis enhances the anti-inflammatory and antioxidant effects of methylprednisolone, leading to reduced activation of inflammatory pathways and increased antioxidant responses during the first hours after injury. These results suggest that metabolic interventions such as ketosis may represent a promising complementary strategy to improve early management of spinal cord injury.

## Introduction

Acute spinal cord injury is a highly catastrophic condition, with high morbidity and mortality in most cases (1), caused by motor vehicle accidents and falls in adults, most frequently in the fourth decade of life (2).

The acute neurological injury caused by direct trauma and compression of the spinal cord is known as primary injury, which causes inflammation of multiple tissue elements due to the release of pro-apoptotic proteins (caspase-1, caspase-2, and cytochrome C), immediate inflammation, ischemia due to loss of perfusion, and increased reactive oxygen species (ROS) (3, 4). This is followed by secondary injury, characterized by lipid peroxidation, necrosis, apoptosis, and neuronal atrophy in the injury penumbra (5, 4). In later phases, increased proinflammatory cytokines (IL-1β and IL-18) inhibit brain-derived neurotrophic factor (BDNF-TrkB) signaling, thereby hindering the recovery of neurological function and neuronal plasticity. (4)

In the initial management of trauma, the time elapsed between the trauma and medical-surgical care is critical. Treatment provides the most significant benefits when performed within the first 24 hours, ideally within the first 8 hours (5). Timely treatment seeks to diminish neuronal damage through surgical spinal cord decompression techniques or the use of drugs that reduce the post-traumatic inflammatory process, such as methylprednisolone, which is routinely used but shows adverse effects at high doses (6, 7). To understand the mechanism of repairing the injury, multiple therapies have been studied. Among them, metabolic modifications in various physiological and pathological conditions are a point of controversy and research. One of these is ketosis, which has been studied in different contexts since 1923 (8). Ketosis is characterized by an increase in ketone bodies (acetoacetate, acetone, and β-hydroxybutyrate) resulting from exogenous administration or endogenous production as an alternative energy source, with relevance to its neuroprotective and anti-inflammatory effects (9).

The metabolism of ketone bodies reduces oxidative stress by stimulating the endogenous antioxidant system via nuclear factor erythroid 2-related factor 2 (Nrf2). In turn, the intracellular NAD^+^/NADH ratio increases, promoting electron transport chain function and mitochondrial biogenesis. Additionally, β-Hydroxybutyrate (βHB) is the primary ligand of the hydroxycarboxylic acid receptor type 2 (HCA-2) expressed in macrophages, dendritic cells, and microglia. The anti-inflammatory effect arises from HCA-2 inhibition of NF-κB, which is associated with the production of prostaglandin D2 by cyclooxygenase 1 (9). Additionally, it inhibits the formation of the NLRP3 inflammasome complex, reducing the expression of IL-1β, IL-18, caspase-1 and 2, cytochrome c, and myeloperoxidase (10). Neuroprotection results from stabilization of the neuronal membrane, leading to an intracellular increase in ATP, a reduction in ROS and lipid peroxidation, stimulation of mitochondrial biogenesis, and modifications in the metabolism of neurotransmitters such as glutamate and GABA, thereby promoting a neuronal survival environment (9). These effects have been described in diseases such as Alzheimer’s, Parkinson’s, and amyotrophic lateral sclerosis (11). Likewise, multiple studies have reported a reduction in the initial lesion area in a ketogenic state, due to increased levels of antioxidant enzymes such as SOD, catalase, glutathione, and Cu-Zn (4).

Another pathway involved in SCI is the hypoxia-inducible transcription factor 1 (HIF-1), which regulates NF-κB gene expression, activating TNF-α and IL-1β. At the tissue level, the increase in these proinflammatory molecules causes oligodendrocyte death by promoting glutamate release, resulting in an excitotoxic environment high in calcium levels (12). The administration of modulators or inhibitors of HIF-1 expression attenuates the secondary inflammatory response and ischemia-reperfusion injury (13, 14, 15). Besides that, the vascular endothelial growth factor (VEGF), down regulated by HIF-1, provokes microvascular disruption, enhances vascular permeability, ionic imbalance, formation of reactive oxygen species, and mitochondrial dysfunction (16, 17) and the histology showed that spinal cord tissue by acute trauma, presented local edema, neuronal death of the gray matter, increased of infarcted areas, and neutrophil infiltration (18).

The establishment of experimental models is limited by the strict dietary regimen required to achieve adequate ketogenic levels. As an alternative, exogenous ketone sources have been used to standardize this process. The use of monoesters at doses of 357–714 mg/kg allows ketone levels to be reached within 30 min, levels only observed on fasting days. The monoester bond is activated in the intestinal wall, generating β-hydroxybutyrate and butanediol, which are absorbed by the liver for the synthesis of new ketone bodies. By this means, measurable ketones are produced in approximately 3 to 4 hours in a dose-dependent manner (19). The purpose of this work is to determine whether exogenous administration or endogenous production of ketone bodies can be achieved through their anti-inflammatory, antioxidant, and neuroprotective effects. An attractive alternative is exogenous ketosis, which is achieved through the administration of βHB precursors. can induce ketosis within 120 minutes of ingestion in a dose-dependent manner (20, 21).

## Materials and Methods

### “Ethics Statement”

Sixty clinically healthy male Wistar rats weighing 250 to 300 g were used. This study was carried out in accordance with recommendations in the NOM 062-ZOO-1999, which refers to the use and care of laboratory animals and the protocol was previously approved by The Institutional Committee for the Care and Use of Animals (IACUC) of Instituto Nacional de Ciencias Médicas y Nutrición Salvador Zubirán (INCMNSZ) (Protocol *number*: **CEX-2024-20/21-1**). All surgery procedures were performed under acepromazine-ketamine as anesthesia. Meloxicam was administrated for analgesia and sodium pentobarbital for euthanasia, and all efforts were made to minimize suffering.

#### Surgical Technique

The animals were anesthetized with acepromazine-ketamine (5-10 mg/kg) intraperitoneally. Meloxicam (1 mg/kg SC) was injected as an analgesic, and enrofloxacin (10 mg/kg IM) was administered. The rat was then intubated orotracheally with a 16G catheter and connected to a rodent ventilator (Kent Scientific RSP 1002). A tidal volume of 6–8 ml/kg, positive end-expiratory pressure of 2–2.5 cm H2O, and a respiratory rate of 60 breaths per minute were maintained. The rat was connected to a rodent anesthesia machine with isoflurane at 2–2.5% minimum alveolar concentration (MAC).

With the rat in the prone position and limb immobilization, a trichotomy of the thoracolumbar vertebral region (T5–L2) was performed and then treated with povidone-iodine embroiled. Next, a longitudinal incision was made on the back of the rat in layers until the thoracolumbar fascia was reached. The paravertebral space was dissected subperiosteally until reaching the spinolaminar space and subsequently up to the ipsilateral transverse process. Hemostasis was maintained at all times. With the support of a surgical microscope (Carl Zeiss, OPMI-1), a laminectomy was performed at the level of T8-T11. Extradural spinal cord clipping was performed using a vascular clip with pressure at 30 grams, and it was removed after one minute of compression (22). Hemostasis was performed. Thoracolumbar fascia was sutured with 2-0 absorbable suture (Vycril) and skin with a continuous suture with 3-0 non-absorbable monofilament suture (Prolene). The experimental subject was placed with water and food *ad libitum* with adequate light exposure to allow physiological sleep-wake cycles. The animals were assisted with manual bladder drainage using abdominopelvic compression.

#### Fasting Time and Dose of 1-3 Hydroxybutanediol monoester

The fasting period was determined at 24 hours, as ketone body levels have been shown to increase at 2.5 mM, and β-hydroxybutyrate levels also increase during this period (23, 24). The dose of the monoester reagent was 357 mg/Kg. At this dose, levels increased after 2 hours and remained there for 5 hours (25).

Group 1 (n=12): Control (CTR). Laminectomy was performed, and intact spinal cord tissue was immediately obtained.

Group 2 (n=12): Spinal cord injury (SCI). Laminectomy and clamping were performed. The animals were euthanized 8 hours after the spinal cord injury.

Group 3 (n=12): Endogenous ketosis (Endo-K+MP). Rats were fasted for 24 hours before spinal cord injury. Laminectomy and spinal cord clamping were performed. Two hours later, methylprednisolone sodium succinate 30 mg/kg IV was administered. They were euthanized 8 hours after spinal cord injury.

Group 4 (n=12): Exogenous ketosis (Exo-K+MP). Monoester was administered at a dose of 357 mg/kg. Laminectomy and spinal cord clamping were performed. Two hours later, 1-3 hydroxybutanediol was administered along with methylprednisolone sodium succinate 30 mg/kg IV, and 5 hours after spinal cord injury, only 1-3 hydroxybutanediol was administered. They were euthanized 8 hours after spinal cord injury.

Group 5 (n=12): Methylprednisolone (MP) (26). A laminectomy and spinal cord clamping were performed. 2 hours later, methylprednisolone sodium succinate (MS) 30 mg/kg IV was administered. The rats were euthanized 8 hours after the spinal cord injury.

After the study period, the animals were anesthetized with sodium pentobarbital overdose (50 mg/kg; IP). The laminectomy incision was then reopened from T8 to T11. The spinal cord was removed. Half of the spinal cord tissue for the histopathological study was preserved in 4% paraformaldehyde immediately after collection. The other half of each group (n=6) was frozen at -70°C for molecular biology studies.

β-hydroxybutyrate levels were determined in each rat using capillary blood collected by caudal puncture with an Abbot apparatus to measure the level of ketosis, induced by fasting or treatment (27).

#### Immunohistochemistry

5 µm slices of each tissue were deparaffinized using xylene for 20 minutes, the tissue was hydrated with ethanol taking it from decreasing concentration of 100% to 50%, washes were performed with phosphate-buffered saline (PBS) to expose the antigens the slides were placed in sodium citrate (10 mM pH6.0) in a water bath for 10 minutes. The peroxidase blocker and primary antibody were placed after washing with PBS. Using the Bio SB kit (BSB 0257, CA, USA), we determined protein expression according to the manufacturer’s instructions. Primary antibodies were probed with following concentrations, HIF-1α 1:500 (SC-13515, Santa Cruz Biotechnology, Ca), NF-κB 1:1000 (SC-8414, Santa Cruz Biotechnology, Ca), NLRP3 1:1000 (SC-134306, Santa Cruz Biotechnology, Ca), IL-1β 1:500 (SC 52012, Santa Cruz Biotechnology, Ca), 3-NT 1:500 (SC-32757, Santa Cruz Biotechnology, Ca), and TNF-α 1:500 (SC-52746, Santa Cruz Biotechnology, Ca). Five microphotographs of each tissue were taken in light microscopy in a LEICA DM 1000 microscope (Leica Microsystems) at 40X magnification. Protein expression was quantified using ImageJ software, with 5 photographs per tissue. A total of 90 microphotographs per group were analyzed to obtain a semi-quantifiable value of peroxidase staining intensity in the spinal cord expressed as a percentage of staining

#### Real-Time PCR

The total RNA was isolated from 50mg of each tissue sample of all groups using TRIzol Reagent (Invitrogen, California, USA). The integrity of the RNA was checked on a 1% agarose gel. First, a reaction with reverse transcriptase (RT) was carried out using 2.5 μg of total RNA in the presence of 200 U of reverse transcriptase (RT) from the Moloney Murine Leukemia Virus (MMLV) (Invitrogen, USA). The amount of mRNA was quantified by real-time PCR on the LightCycler 480 Roche through the reaction with SYBR Green. The method to calculate the amount of mRNA was assessed by ΔCT (increase of the cycle threshold) for (NLRP3), and the sequences used were the following: 5’-GCTGCTCAGCTCTGACCTCT-3’ and 5’-AGGTGAGGCTGCAGTTGTCT-3’, (HIF-1α) 5’-GTAGTGCTGATCCCTGCACTGAATC-3’ and 5’-CTGGGACTGTTAGGCTCAGGTG-3’, (NF-κB) 5’-TAAAAGCTGGCAAGCGTATCC-3’ and 5’-CCTGCTGCCGCTCATTACA-3’, (IL-1β) 5’-GGGATTTTGTCGTTGCTTGT-3’ and 5’-CAGGAAGGCAGTGTCACTCA-3’, (TNF-α) 5’-ATGAGCACGGAAAGCATGAT-3’ and 5’-CTCTTCAAGGGACAAGGCTG-3’, (VEGF) 5’-CTTGGGTGCACTGGACCCT-3’ and 5’-CACTTCATGGGCTTTCTGCTC-3’. The levels of mRNA were normalized using a GADPH control (5’-GTCATCATCTCCGCCCTTCC-3’, 5’-CACGGAAGGCCATGCCAGTGA-3’) (28).

#### Statistical analysis

Results were expressed as mean ± standard error. Inferential statistics were used to compare means between groups using an ANOVA test and a Tukey post-hoc test, using PRISMA software. A p-value <0.05 was considered statistically significant.

## Results

Wistar rats weighing 250-300 g were used, and the groups did not show any weight difference. Before surgical procedures, ketone bodies were measured to confirm their increase, as shown in Figure 1. There was a statistically significant difference showing that End-k+MP and Exo-k+MP had higher levels of ketosis than CTR, SCI, and MP.

**Figure 1.** Blood Ketone, End-k+MP, Exo.k+MP *P<0.05 vs CTR, SCI, MP

Ketone Bodies are normally produced as an alternative source of energy in times of fasting. The human body, via ketogenesis, produces acetone, acetoacetate, and beta-hydroxybutyrate (29). It has been shown that this “ketosis state” modulates inflammation, interleukins, and oxidative stress. We decided to specifically address 2 pathways: oxidative stress and the inflammation pathway.

ROS formation is a crucial component of secondary injury, exacerbating damage and contributing to a poor prognosis. Nrf2 is a transcription factor that regulates the expression of antioxidant genes; our results showed that Nrf2 levels may be reduced after SCI, but these are increased by MP and synergistically when ketosis, either Endo or Exo, is used alongside, therefore increasing the antioxidant response (Figure 2). There are no studies regarding glucocorticoids and activation of Nrf2, but it seems there is no direct bond. An increase in Nrf2, as shown in our data, may be because of the anti-inflammatory response, leading indirectly to an activation of the transcription factor. Our data also showed an increase in protein nitration and, therefore, oxidative stress, using 3 Nitrotyrosine (3NT) as its marker. Our results (Figure 3) showed that all treatments had a significant difference against SCI, indicating their effectiveness. Nevertheless, Endo-k worked again synergistically with MP, having a greater impact than MP alone, the traditional treatment.

**Figure 2.**
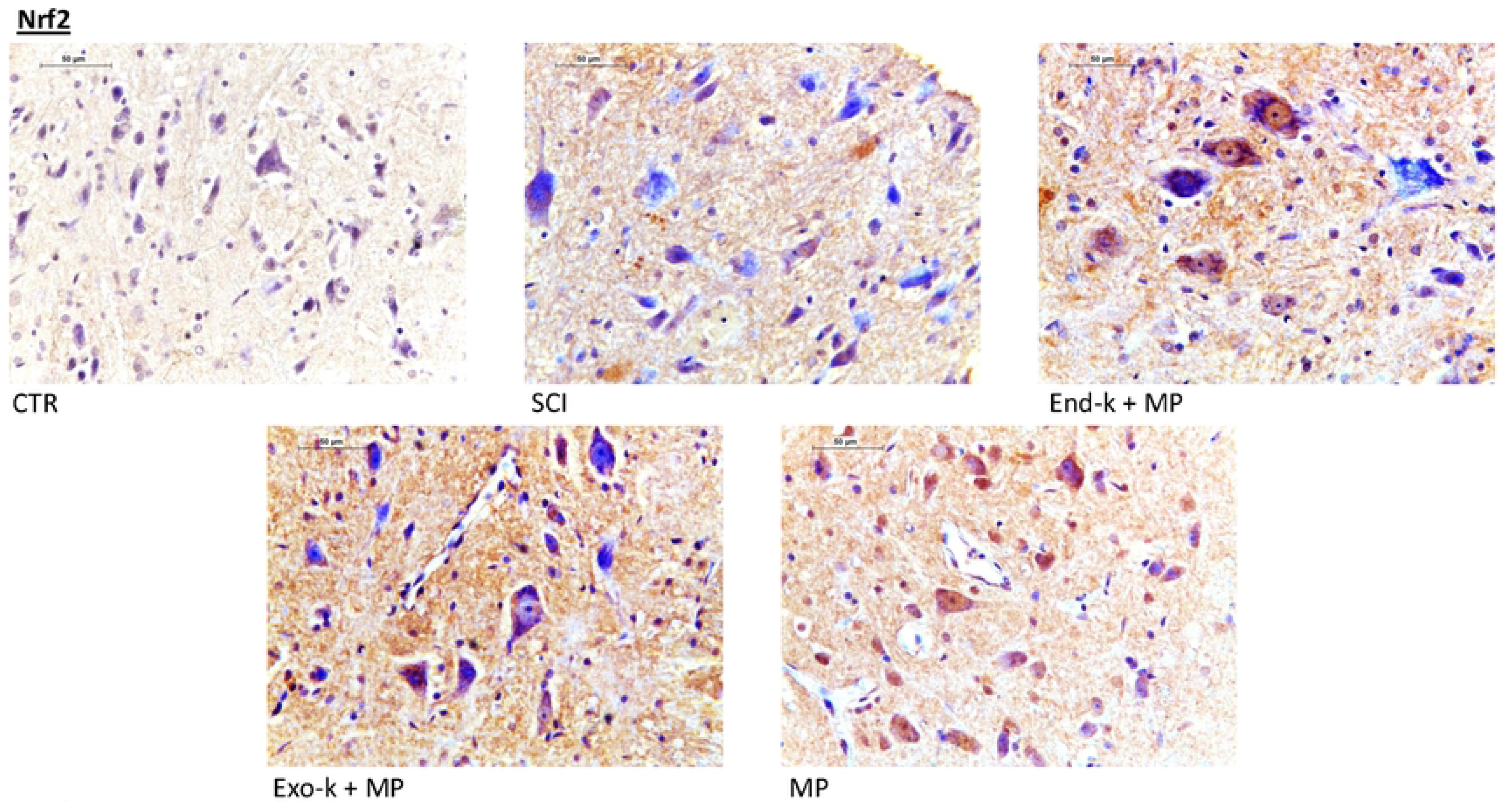
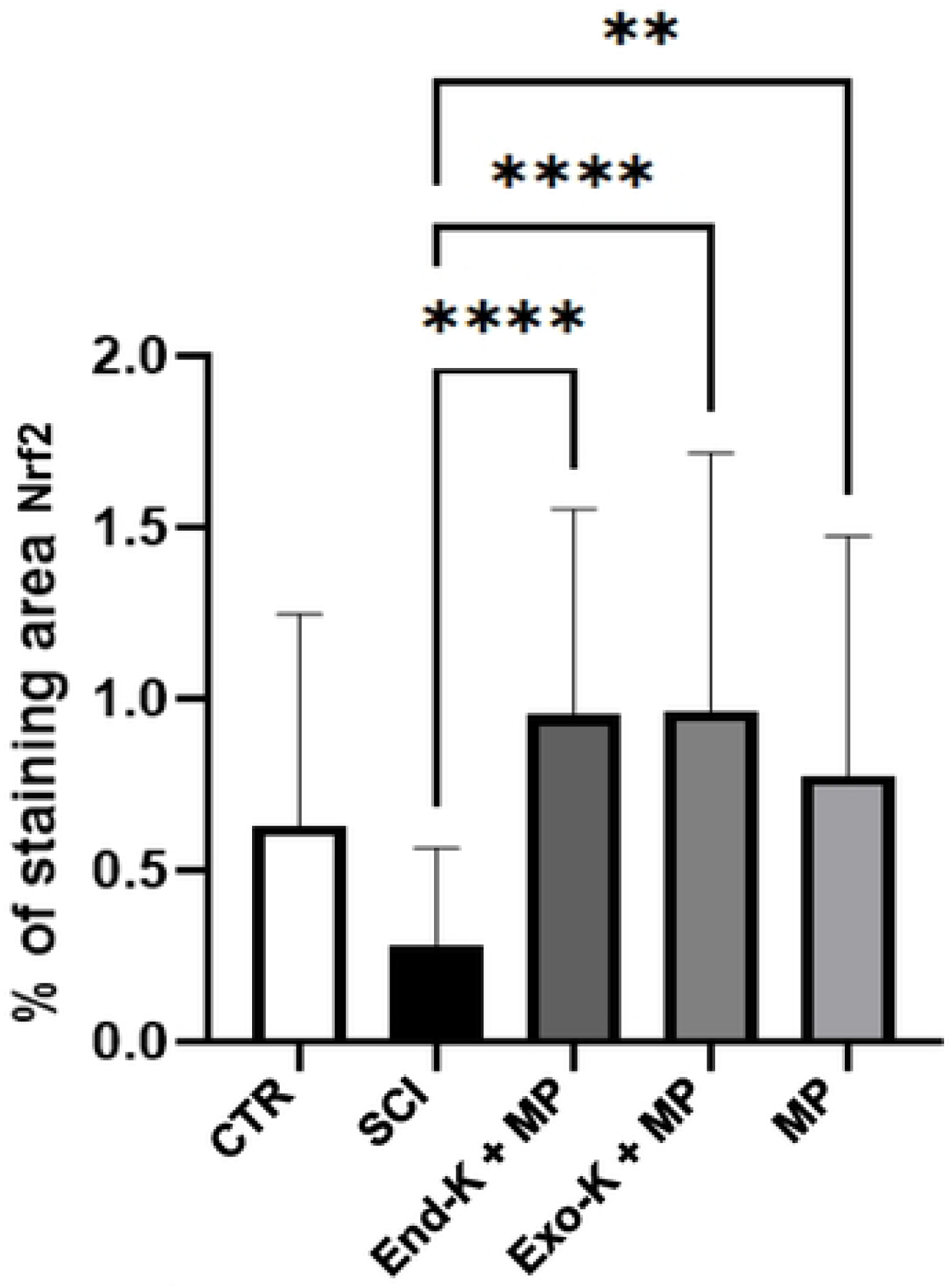
a) Immunohistochemistry (X40) of spinal cord tissue b) Nrf2 was expressed as mean ± S.E., groups were n=6, ****P<0.05 vs Endo-k+MP, Exo-k+MP, **P<0.05 vs MP

**Figure 3.**
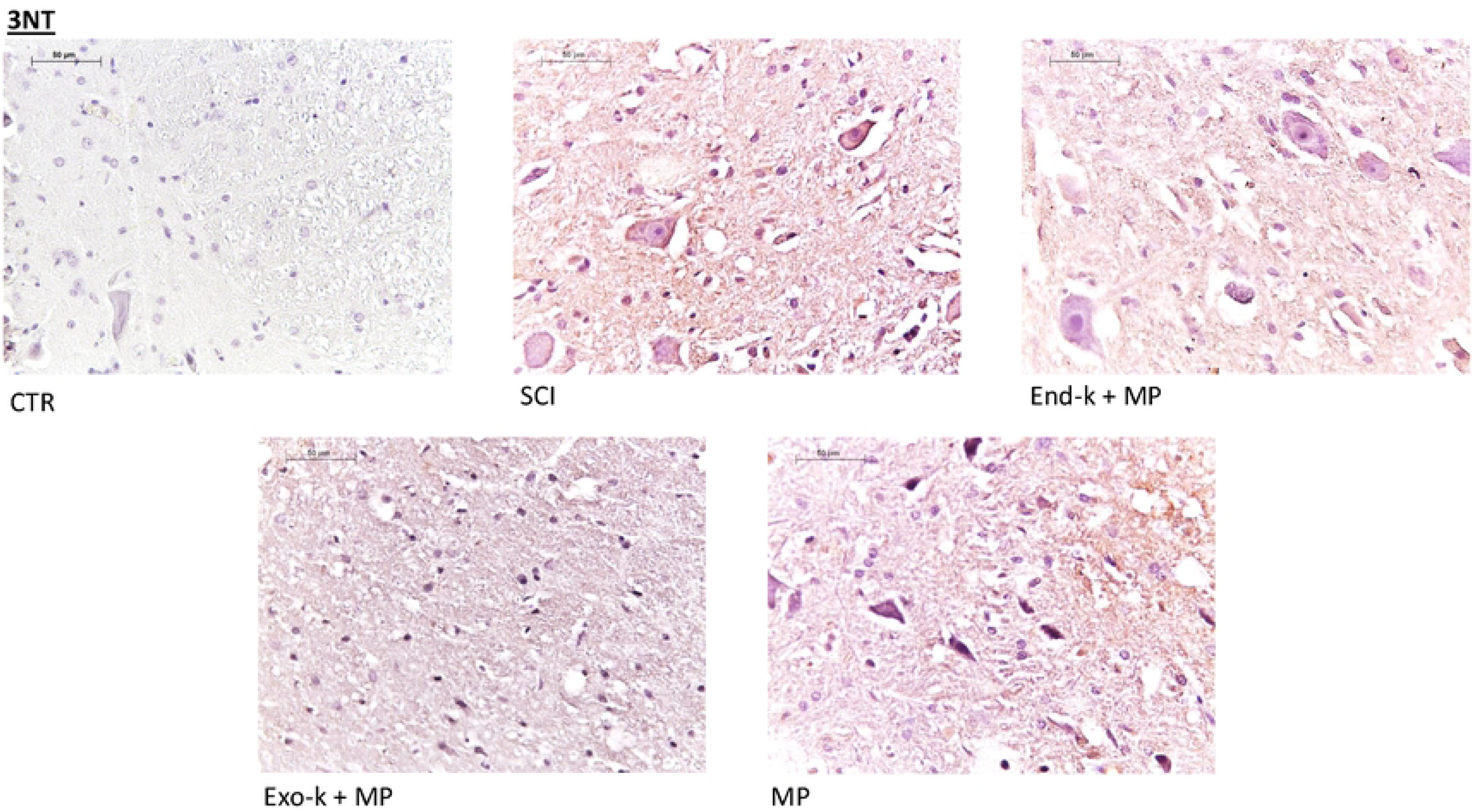
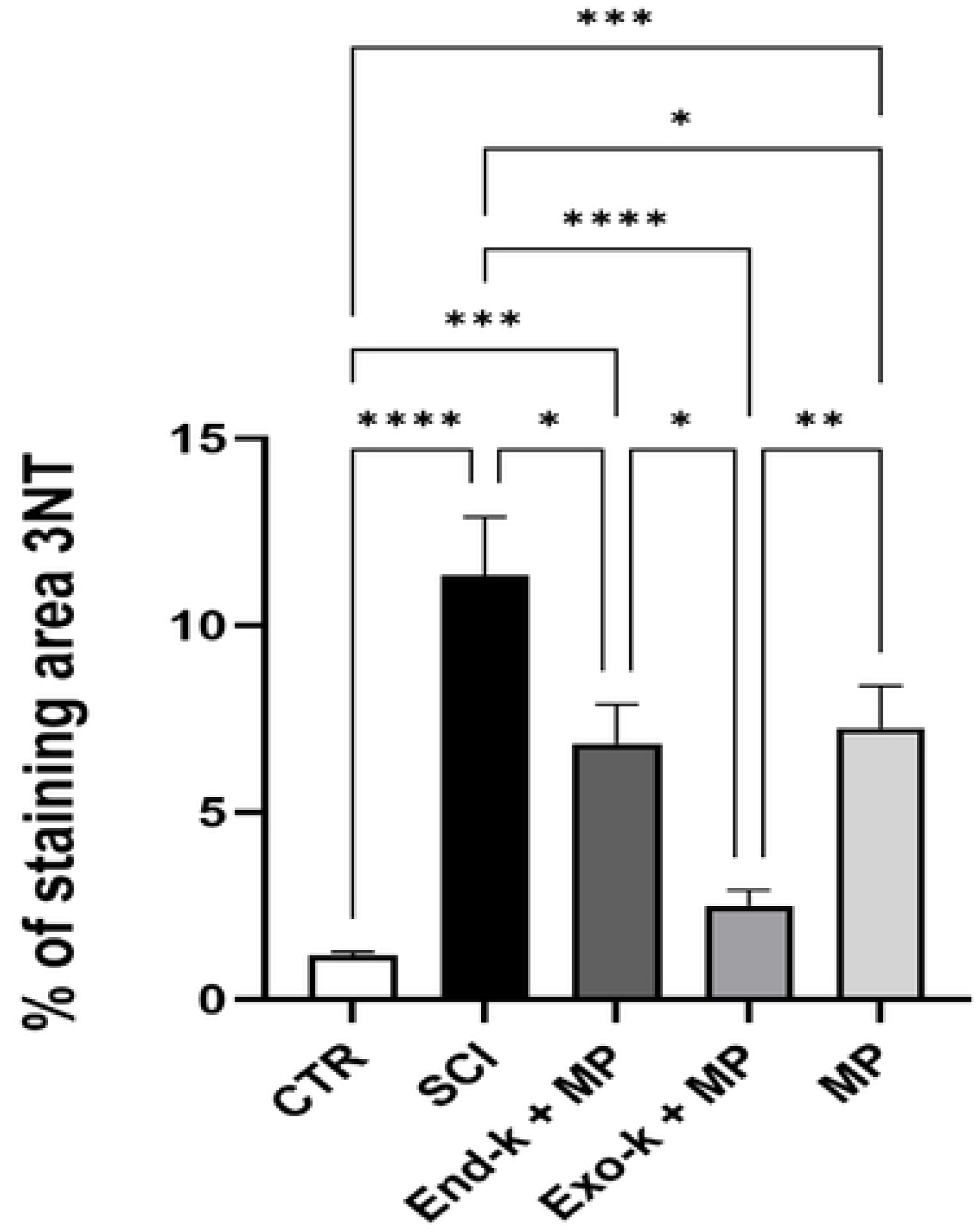
a) Immunohistochemistry (40X) of 3NT as an oxidative stress marker b) 3NT expression all results are expressed by mean ± SE, *p<0.05: SCI End-k+MP, Exo-k+MP, MP ** p<0.05 vs Exo-k+MP, MP. ***p<0.05 vs CTR, End-k+MP, MP. ****p<0.05 vs CTR, SCI, Exo-k, MP.

Furthermore, the inflammatory response is also affected by ketone bodies, as described previously. They directly inhibit HIF-1 and NF-κB, and the inhibition of these factors also causes a chain reaction that leads to the downregulation of proinflammatory chemokines, such as TNF-α and IL-1β. Additionally, NLRP3 formation is also inhibited, preventing the release of IL-1β and IL-18, two proinflammatory cytokines.

HIF-1 is triggered by hypoxia, ROS, inflammation, and several other metabolites; the results of the immunohistochemistry revealed End-k + MP and MP were effective in decreasing levels of the transcription factor. But its levels in Exo-k + MP were equal to the SCI group. In the case of PCR, mRNA levels showed no statistical difference, regardless of the treatment used. (Figure 4)

**Figure 4.**
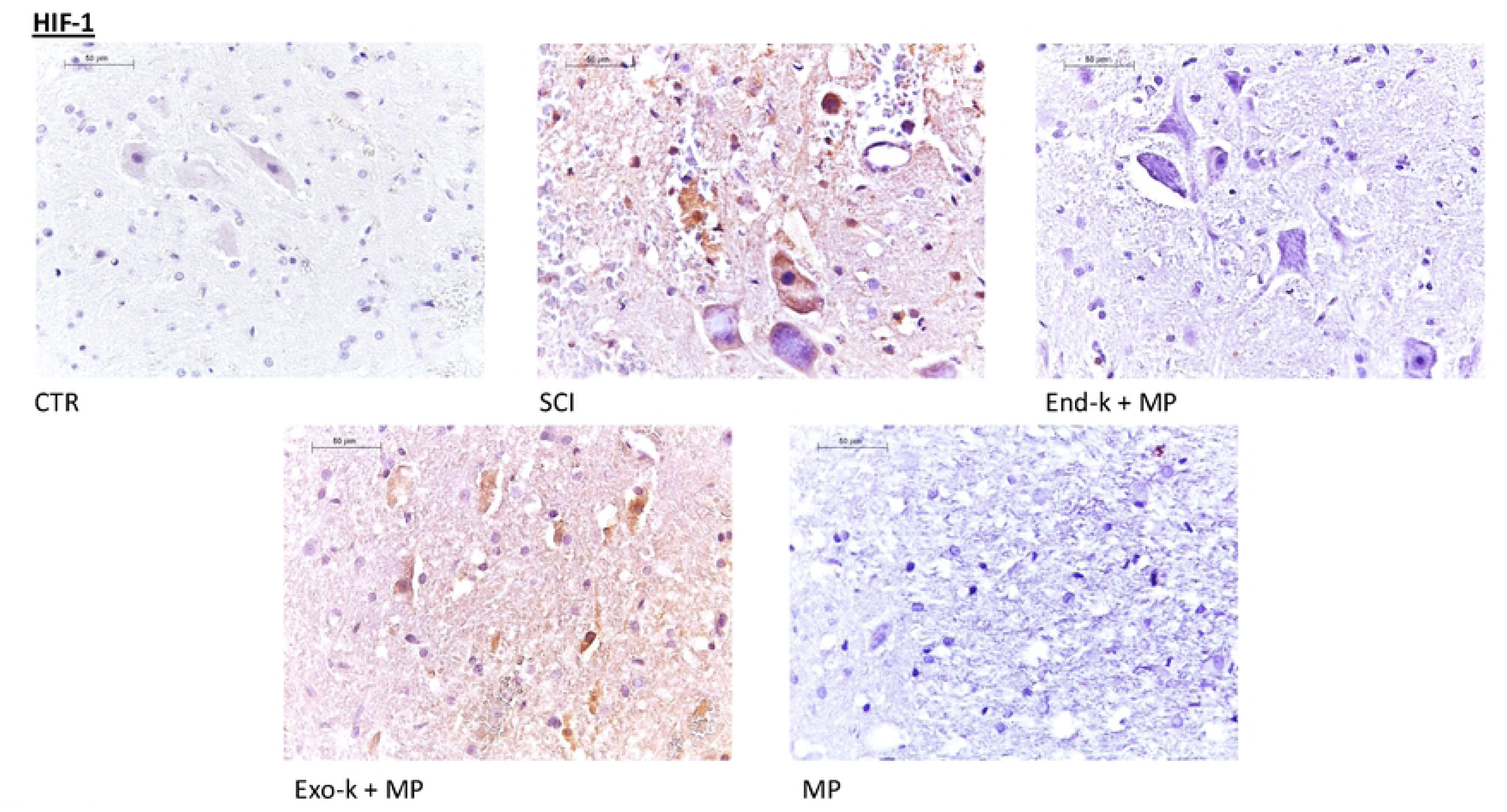
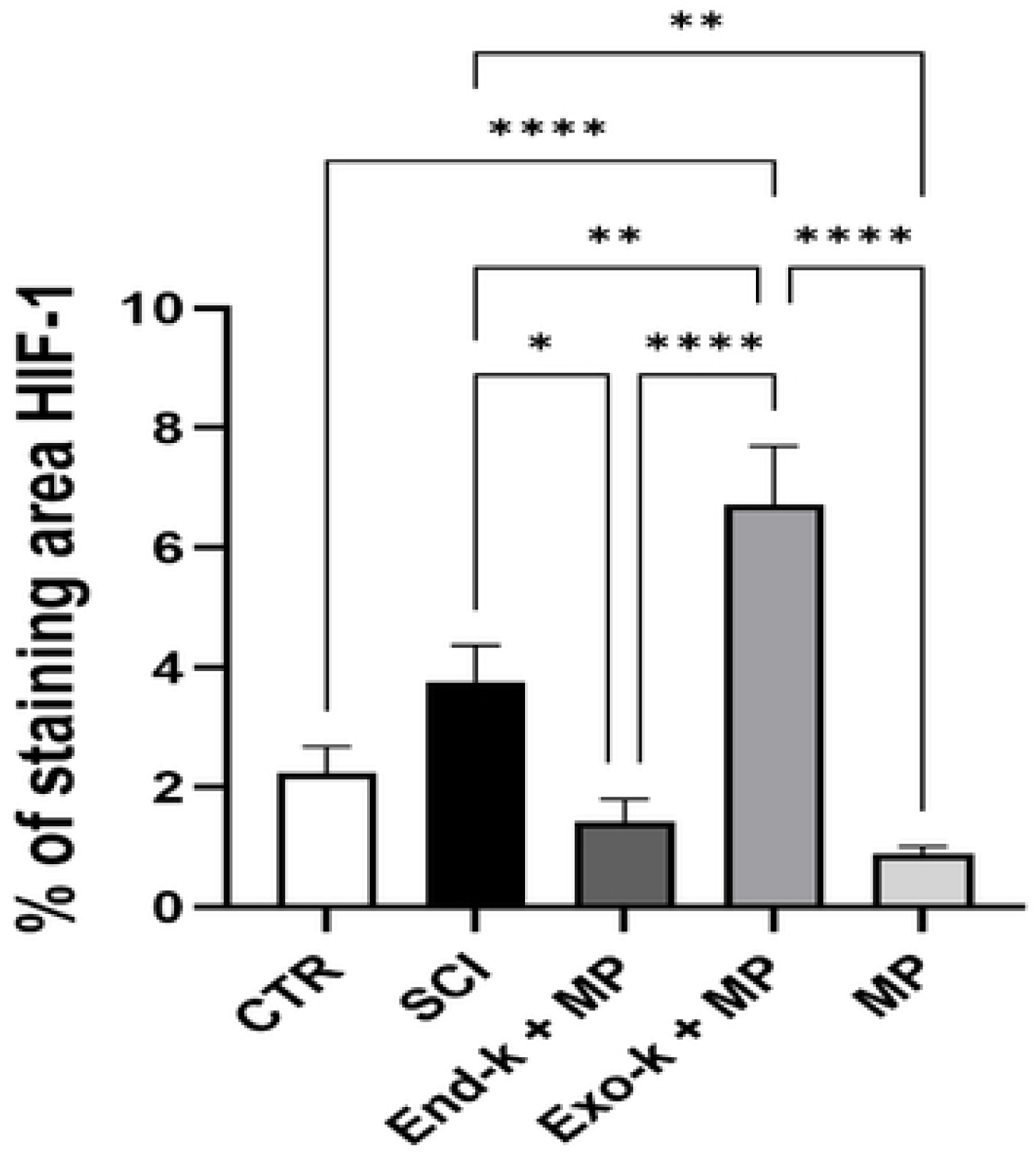
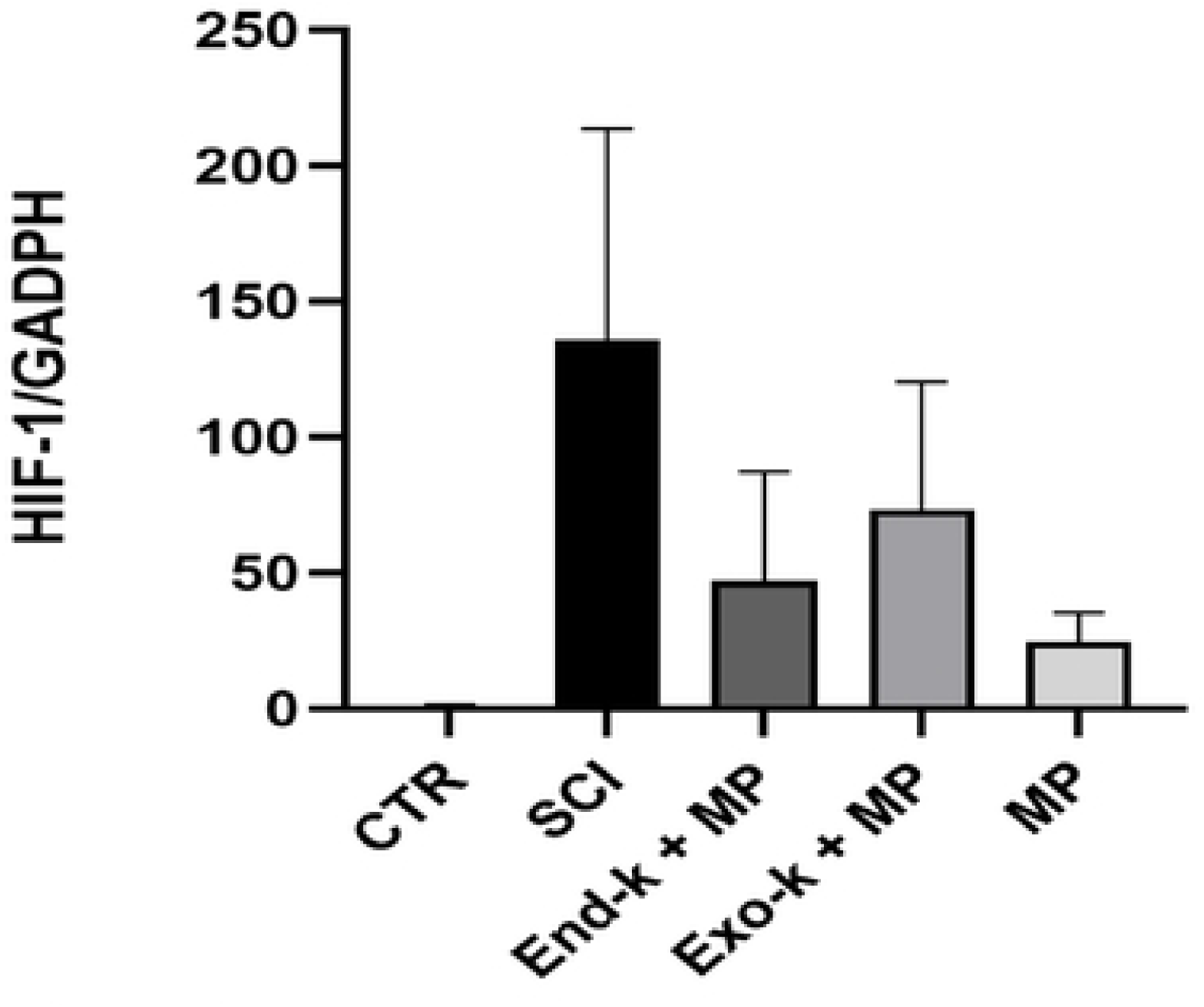
a) Immunohistochemistry (40X) of HIF-1, b) HIF-1 protein expression, all results were expressed in mean ± S.E. *p<0.05 vs SCI, End-k+MP. **p<0.05 vs SCI, MP. **** p<0.05 vs End-k+MP, Exo-k+MP, MP, CTR. c) mRNA levels of HIF-1, and each bar represents the HIF1/GAPDH ratio; there is no difference among treatments.

NFkB is normally inactive in the cytoplasm and activated by proinflammatory chemokines and ROS. In the immunohistochemistry, the data revealed that all treatments had an impact on its activation compared to the SCI group; nevertheless, End-k + MP worked more effectively than Exo-k + MP to the point that End-k + MP was the only treatment that did not have a statistical difference compared to CTR.

Similarly, in the PCR, CTR, and End-k + MP were the only groups with the mRNA levels statistically different from SCI. (Figure 5)

**Figure 5.**
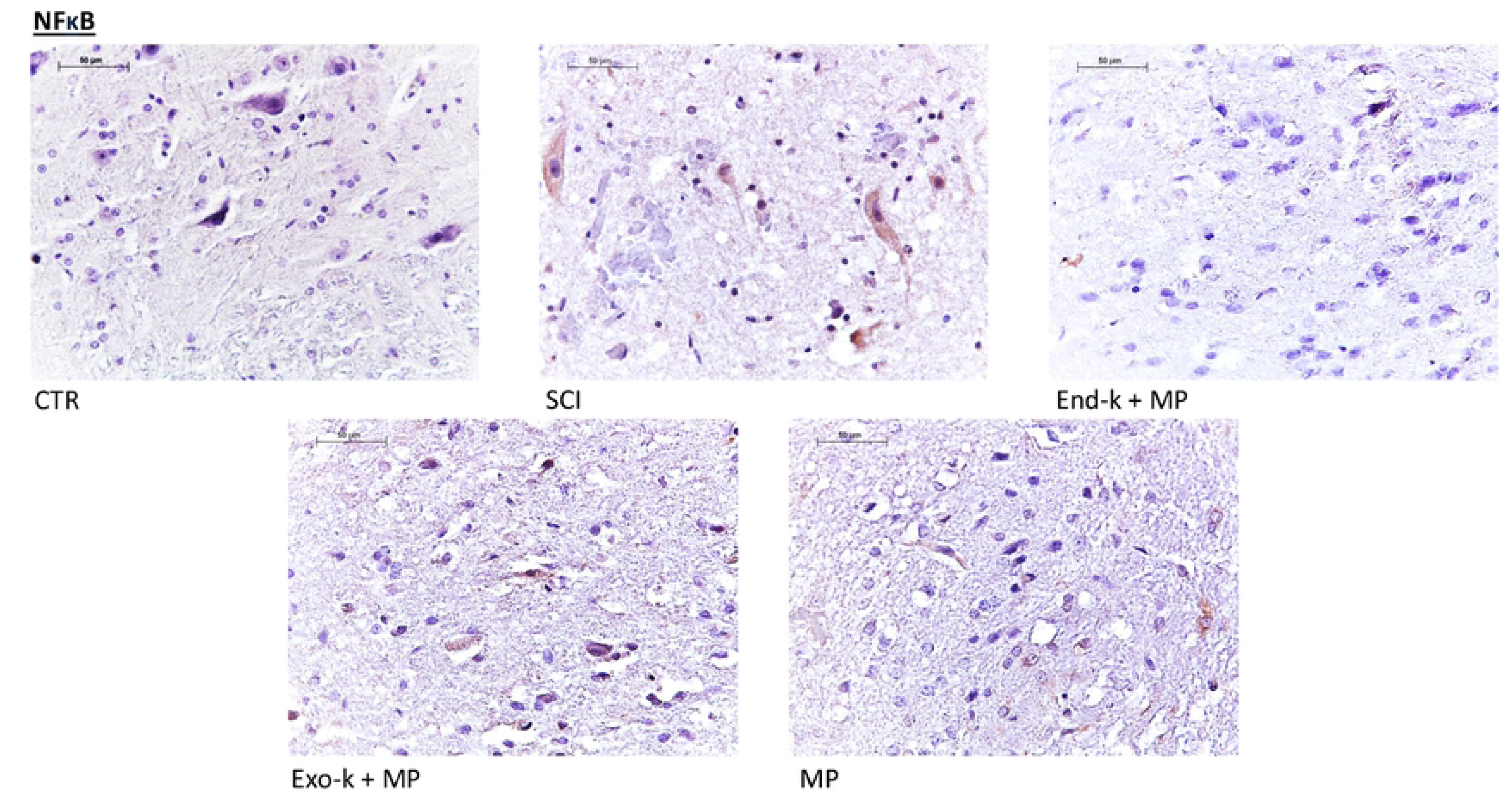
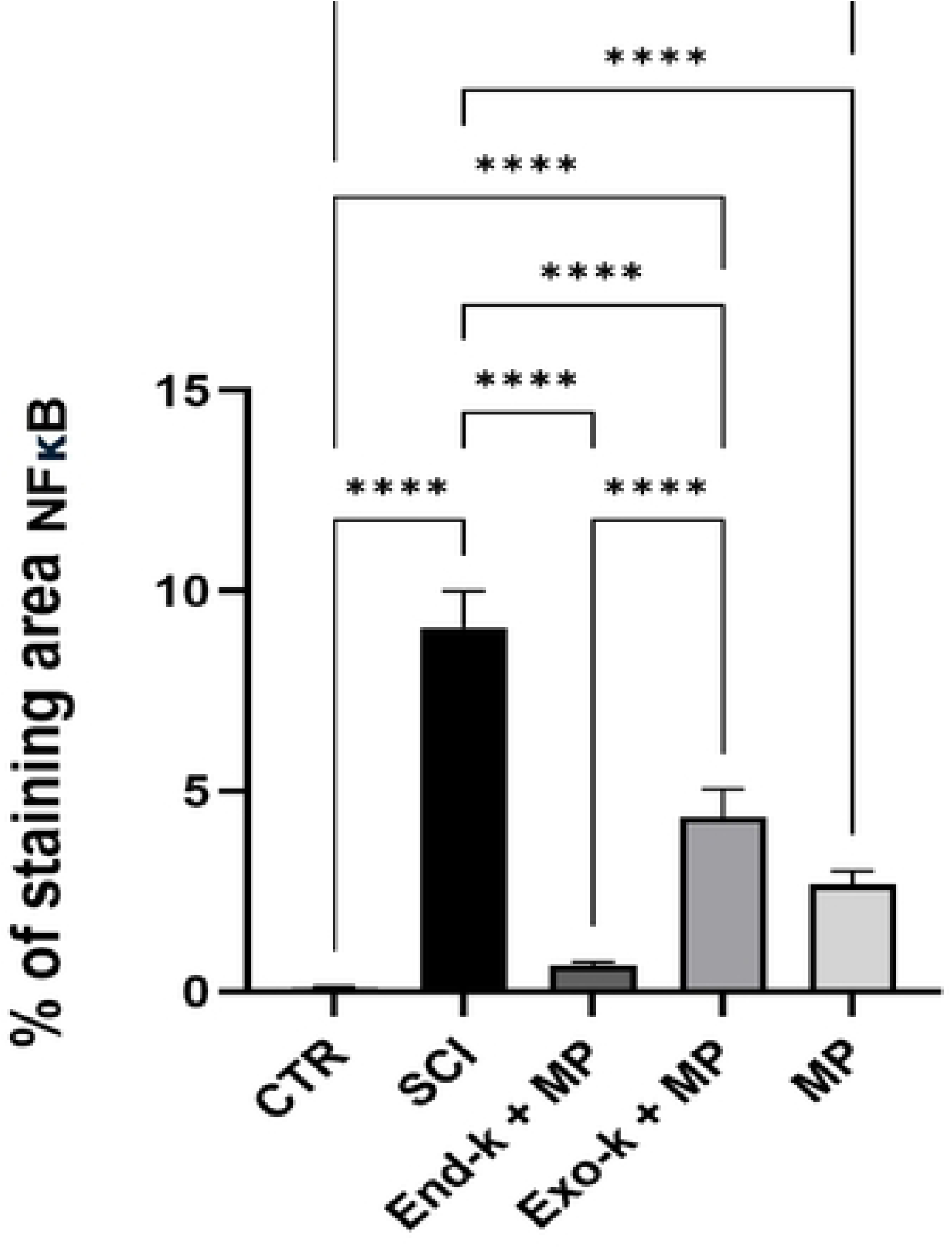
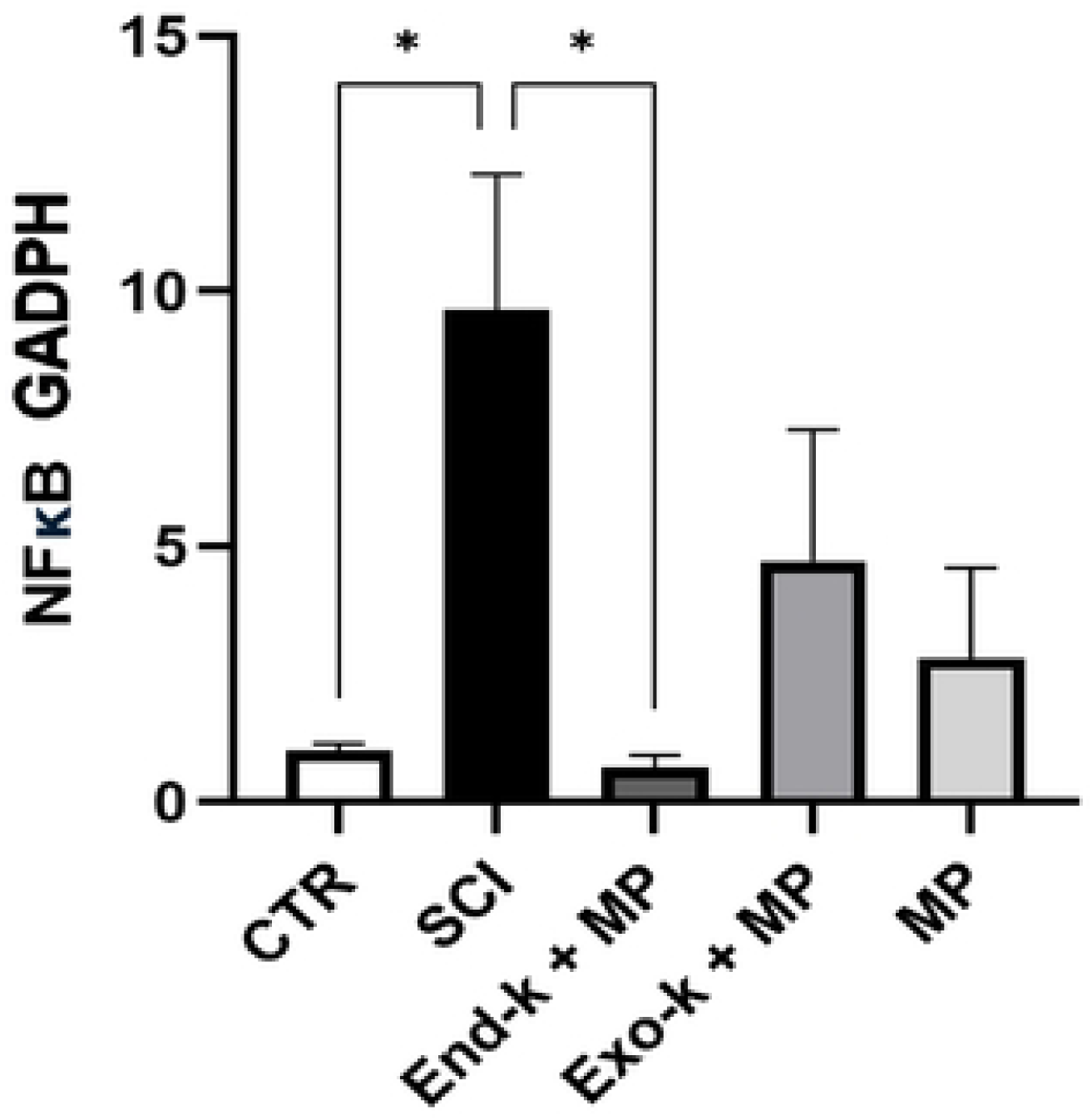
a) Immunohistochemistry (40X) of NFkB b) All results were expressed in mean ± S.E. SCI **** P<0.05 vs CTR, End.k + MP, Exo-k +MP, MP. CTR ** P<0.05 vs MP. End-k + MP P<0.05 vs Exo-k + MP. c) mRNA levels of NFkB. Each bar represents the NFkB/GAPDH ratio, SCI * P<0.05 vs CTR, End-k + MP.

NLRP3 is the inflammasome that activates proinflammatory chemokines and promotes apoptosis. Data in immunohistochemistry revealed a downregulation by every treatment used, with no statistical difference between them or the CTR group. Furthermore, in the PCR, there was no statistical difference among any group. (Figure 6)

**Figure 6.**
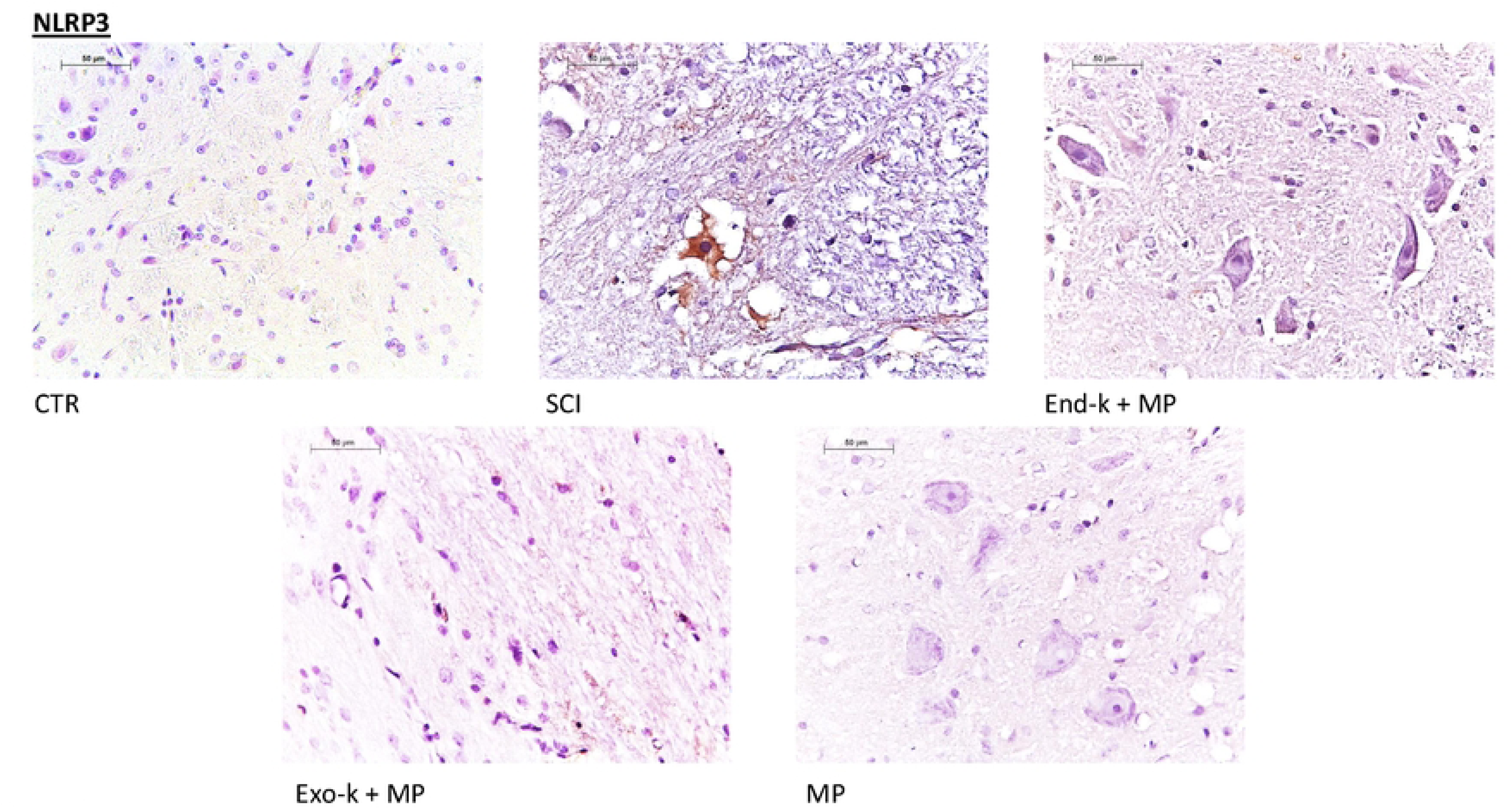
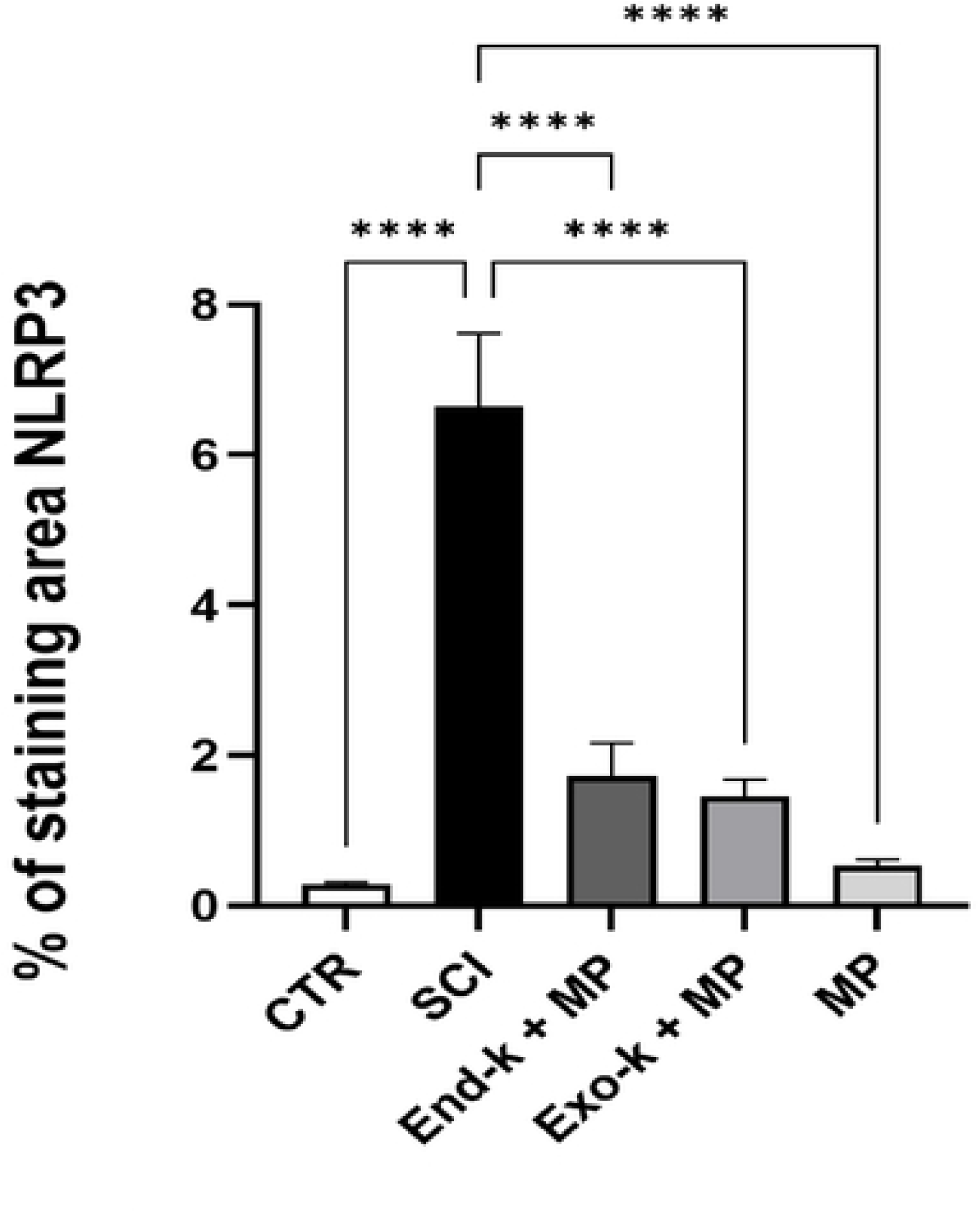
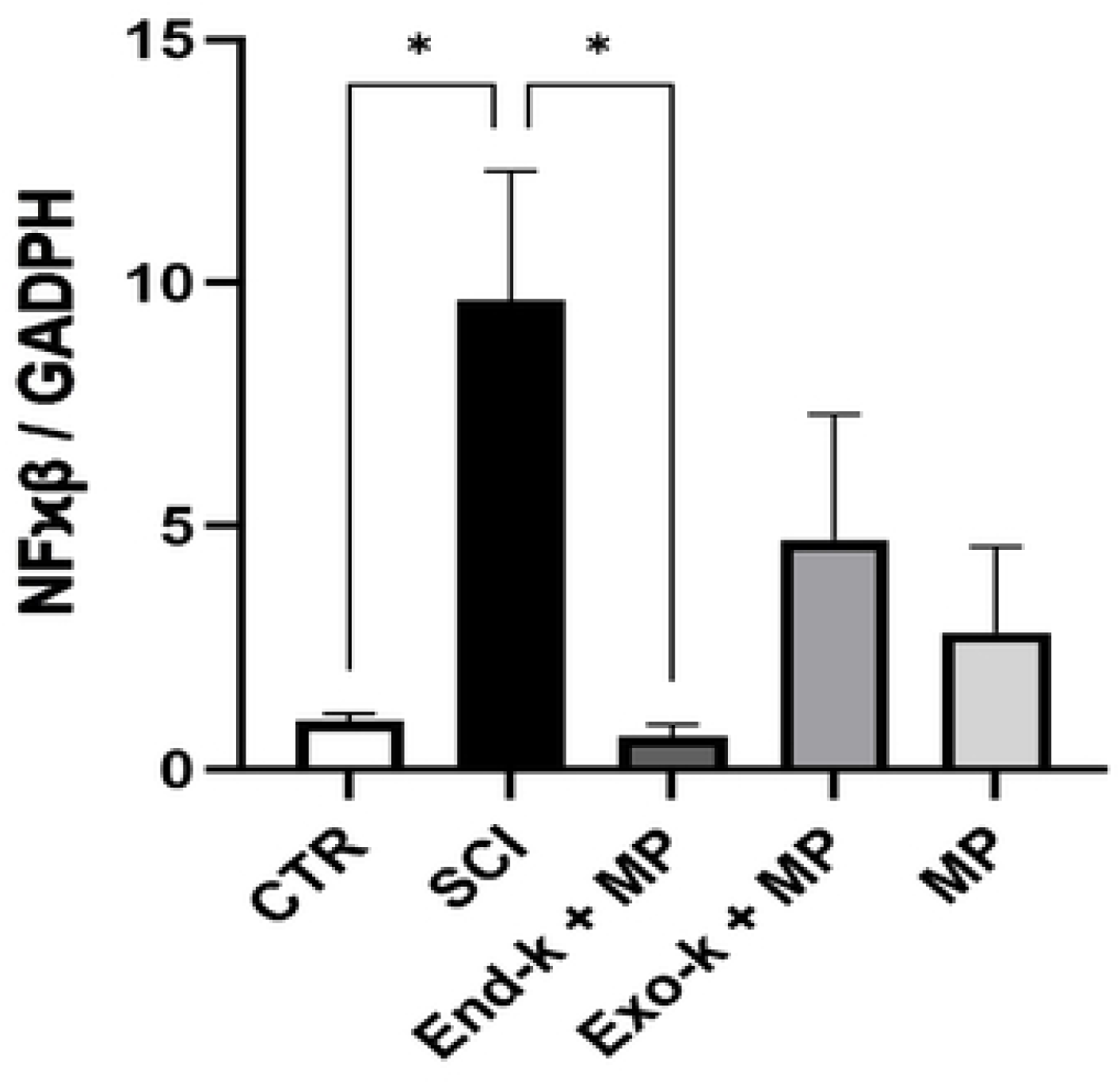
a) immunohistochemistry (40X) of NLRP3 inflammasome, b) the graph shows the levels of NLRP3 protein, all results were expressed in mean ± S.E. ****p<0.05 vs CTR, SCI, End-k+MP, and MP, c) mRNA levels of NLRP3, each bar represents the NLRP3/GADPH ratio, there was no difference among treatments.

TNF-α, a proinflammatory chemokine, is promoted by all transcription factors addressed before, but our data revealed a decreased expression when treatment is used. Exo-k + MP was the only one not statistically different from CTR, indicating that MP and Exogenous ketosis work effectively and synergistically. In the PCR, the mRNA levels did not show a difference. (Figure 7)

**Figure 7.**
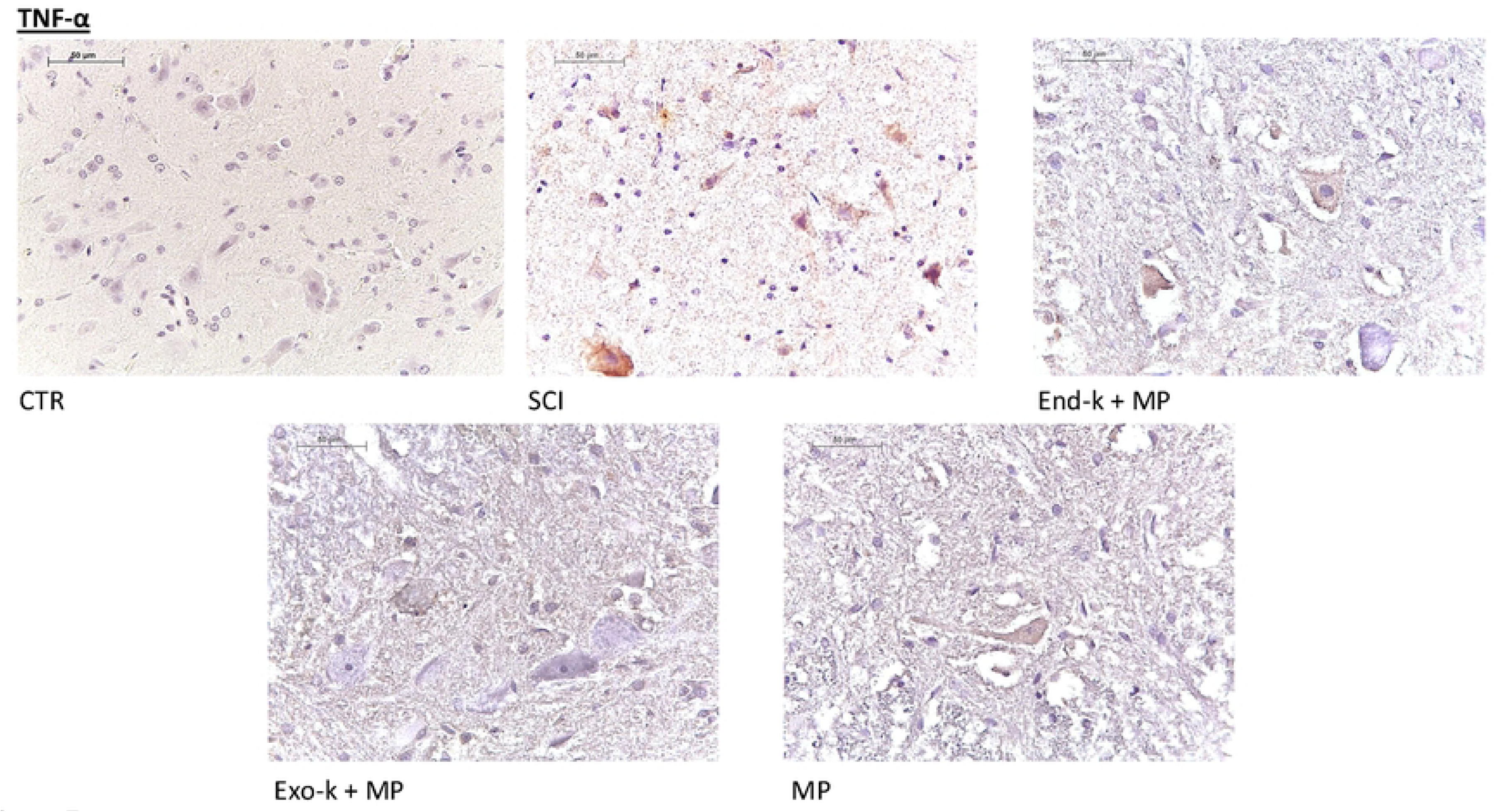
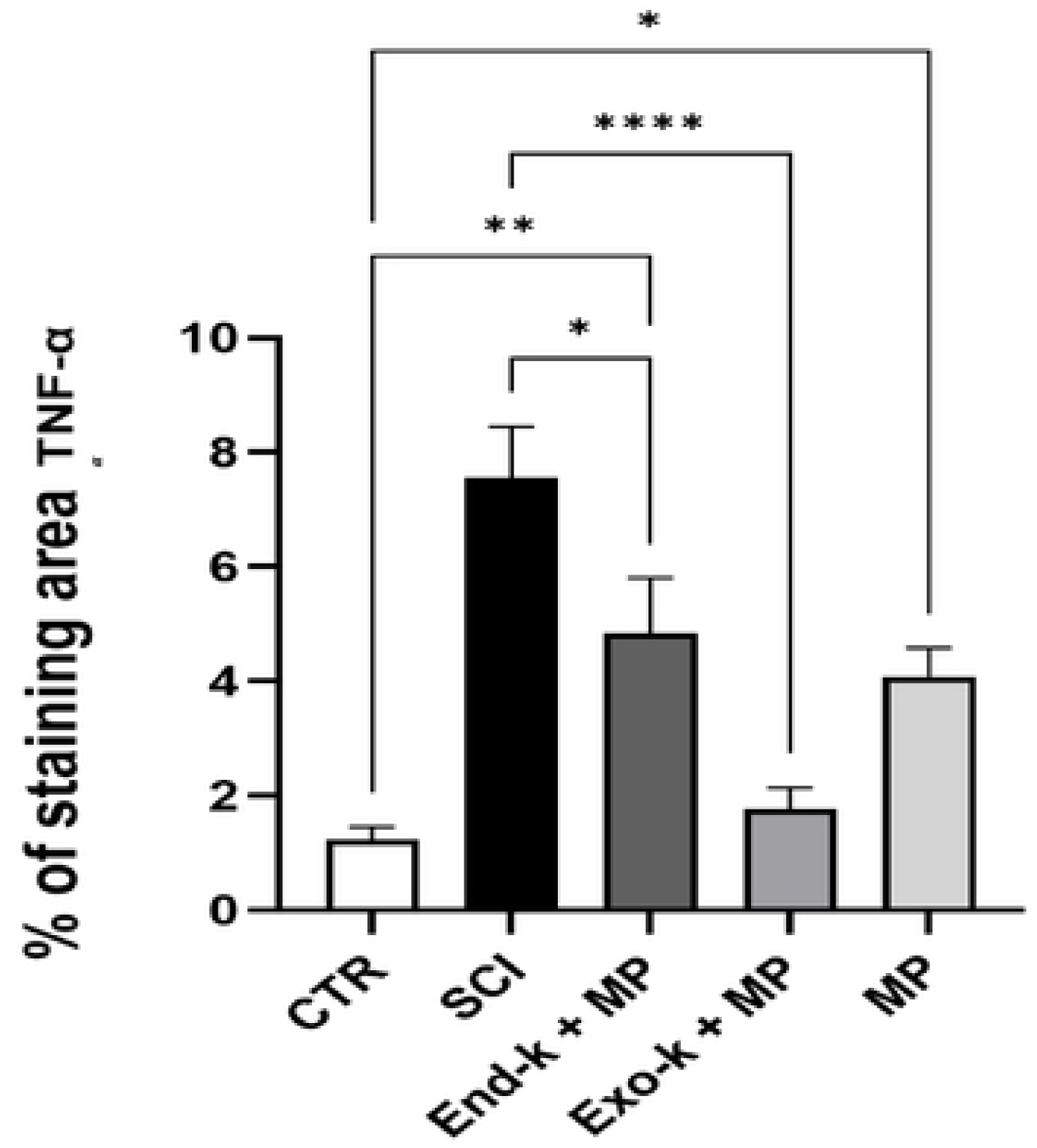
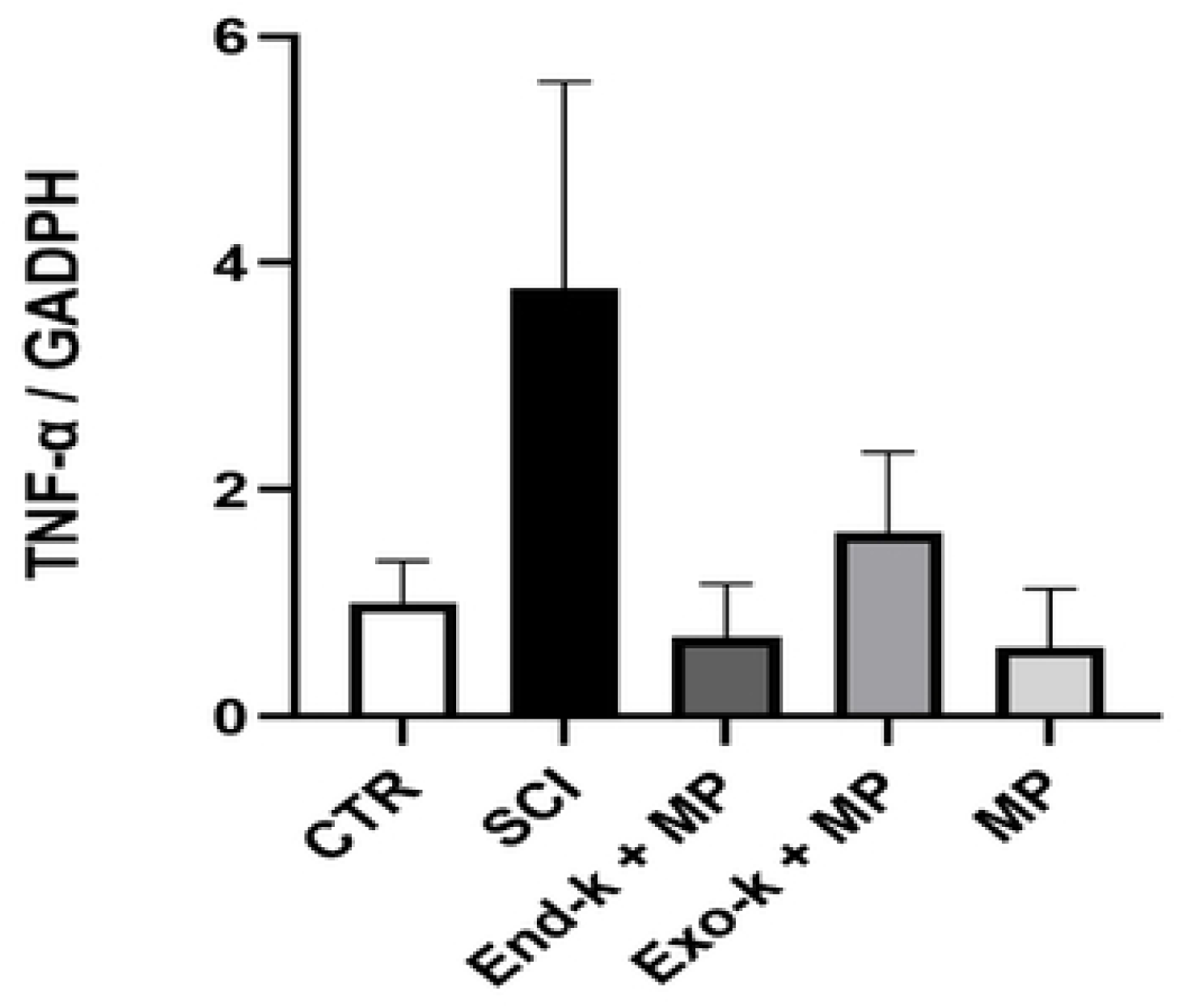
a) immunohistochemistry (40X) of TNF-ɑ. b) All results were expressed in mean ± S.E. SCI **** P<0.05 vs Exo-k + MP, SCI * P<0.05 vs End-k + MP, CTR * P<0.05 vs MP., CTR ** P<0.05 vs End-k + MP.. c) mRNA levels of TNF-ɑ. Each bar represents TNF-ɑ/GAPDH ratio; it showed no difference among treatments.

Cytokine IL-1β is also upregulated by transcription factors and the inflammasome addressed before. In immunohistochemistry, the data revealed that all treatments were effective in downregulating the chemokine, which was in agreement with the PCR results. (Figure 8)

**Figure 8.**
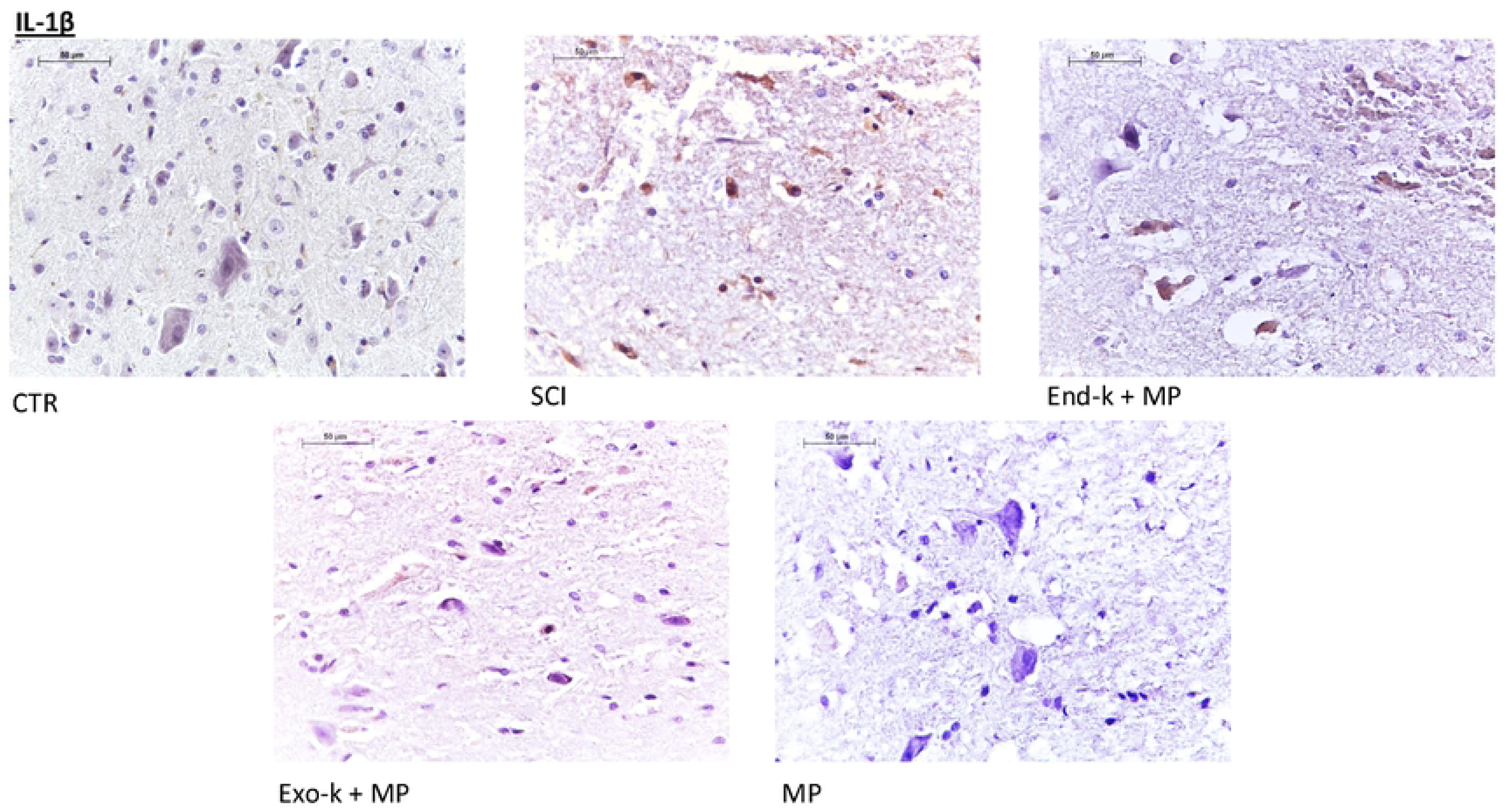
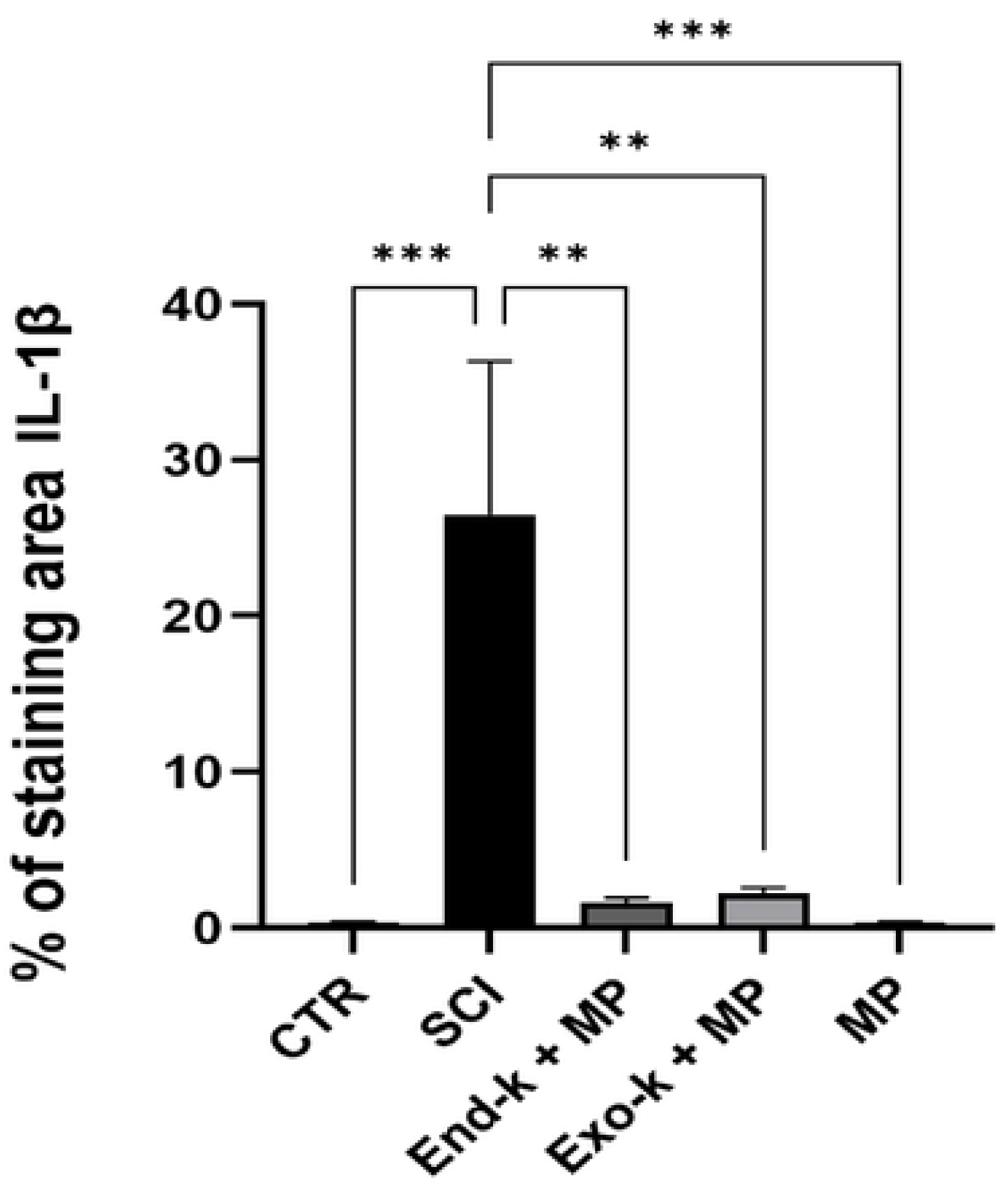
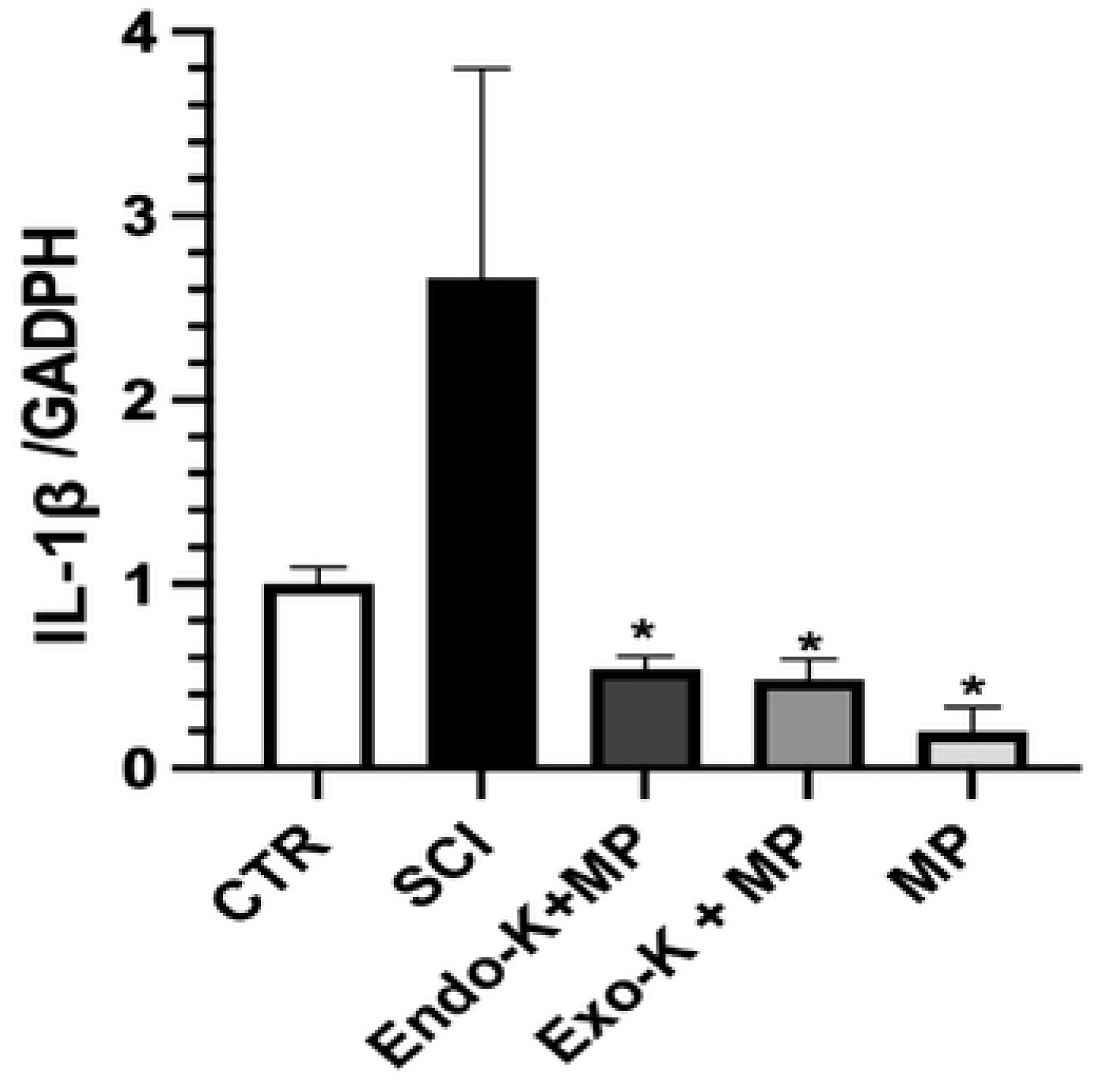
a) Immunohystochemistry (40X) of IL-1β, b) All results were expressed in mean ± S.E. SCI *** P<0.05 vs CTR, MP. SCI ** P<0.05 vs End-k + MP, Eco-k + MP. c) mRNA levels of IL-1β. Each bar represents IL-1β/GAPDH ratio, SCI * P<0.05 vs MP, Exo-k+MP, and End-k+MP.

## Discussion

The incidence of SCI and its impact on the quality of life of patients make it important to research better treatments and ways to mitigate secondary injury. In this study, the first eight hours of acute spinal cord injury were studied. During the secondary injury phase, an inflammatory and hemorrhagic response characterized by ischemic damage and/or apoptotic cell death occurs. Microglia, T cells, and astrocytes are subsequently activated. There is an exacerbated release of cytokines such as IL-1β and TNF-α, as well as an overexpression of transcription factors such as HIF-1α, VEGF, and NFκB.

Several studies have explored the possibility that ketosis regulates the inflammatory response in the central nervous system, and some studies indicate that the ketogenic diet may be beneficial in neurological disorders such as Alzheimer’s disease, Parkinson’s disease, amyotrophic disease, and multiple sclerosis (11). However, the effects of ketone bodies in spinal cord injury are not completely elucidated.

It has been observed that the ketogenic diet improved forelimb motor function after spinal cord injury in rodents. (30, 31)

Ketogenesis is the response to energy demands when there are low levels of carbohydrate, insulin, and high levels of glucagon and cortisol, which is also autoregulated by inhibiting lipolysis through PUMA-G nicotinic receptors in adipocytes and by fatty acid flux, acetyl-CoA concentrations, and enzymatic deacetylation in hepatocytes.

This diversity of autoregulation methods makes it difficult to accurately measure EndoK levels, which facilitates the regulation of concentrations when ExoK is used. This ExoK is released into the portal circulation as βHB and butanediol in equal amounts; (32), the latter is then transformed into βHB by alcohol dehydrogenase in the liver; therefore, increasing bioavailability of βHB, the dose to achieve “endogenous levels” was 357–714 mg/kg (19). Recognizing that ketones have a short half-life (33), this is why ketone levels could vary dramatically depending on whether the subject is calm or under stress.

In this work it is shown that once 1-3 Hydroxybutanediol Monoester was administered, the bioavailability of beta-hydroxybutyrate increases and the levels of superoxidodismutase (SOD) are increased. SOD is an antioxidant through the acetylation of histones and FOXO1 / 3 and Nrf2, in addition to increasing the transcription of Catalase (CAT) and heme oxygenase (HO-1), and through the tricarboxylic acid pathway that uses fumarate at the mitochondrial level. One of the antioxidant effects is that SOD acts as a carrier of O₂⁻ and (•OH). (34) In the absence of O₂⁻, it will not bind to NO, and ultimately, there is no abundant production of peroxynitrite, ONOO⁻. Because of this, the direct effect on the alterations in protein tyrosine residues and intracellular amino acids is diminished during the first 8 hours of acute spinal cord trauma. This is why, when 1-3 Hydroxybutanediol Monoester is administered during the first two hours in rats with acute spinal cord trauma, a decrease in 3-NT is shown, in contrast to treatment with MP and compared to rats that had been fasted for 24 hours.. Furthermore, once ketogenesis is induced through the administration of 1-3 Hydroxybutanediol Monoester, there is an increase in the availability of succinate at the intracellular level (35), which in turn causes an inhibition of prolyl hydroxylases (PHD). Puchowicz M. et.al stimulated rats with a ketogenic diet for 4 weeks and then subjected them to focal cerebral ischemia. In addition, the same ischemic process that the spinal cord tissue is undergoing due to the injury results in HIF-1α not being ubiquitinated, therefore, it is not degraded and, consequently, it is translocated to the nucleus and dimerizes with the HIF-1β subunit, resulting in a synthesis of its HIF-1 protein, as could be observed when 1-3 Hydroxybutanediol Monoester was administered in our work evaluating rats with acute spinal cord trauma during an 8-hour study.

Shippy et al. proved that βHB alleviated the neurodegenerative process in Alzheimer’s disease (AD) in a transgenic mice model by the union of GPR109A and, therefore, inhibiting NF-κB. Luet et al. also described a decrease in NF-κB and an increase in Nrf2 (antioxidant pathway) with the use of ketogenic diet (KD), related to a better response on behavior and electrophysiological recovery (36) and an increase by three times in the concentrations of HIF-1a and Bcl-2. (37, 38)

Additionally, D’Ignazio L. et al. also demonstrate an interaction between hypoxia and inflammation, showing the presence of oxygen-independent transcription factors, such as STAT2 and NFkB. Interestingly, not only does NFkB induce a HIF-1 response, but HIF-1 itself also has NFkB regulatory activity under inflammatory conditions. (39) It should be acknowledged that when SCI occurs, not only inflammation is key part of the secondary injury, but also ischemia (being hipoxia the main signal for HIF-1 to express), vascular damage, apoptotic signalling, excitotoxicity, demyelinitation, fibroglial scar, and far more other mechanisms (40) Using ExoK the way it was used in this study may be not enough to counter all of these mechanisms, and this could explain whyindividuals induced into endogenous ketogenesis showed lower transcription of HIF-1α, as well as NFkB, compared with ExoK induced individuals. This could also be related to an inflammatory response in addition to the HIF-1 stimulation induced by succinate as a substrate under hypoxic conditions suffered by the bone marrow tissue (41).

Reactive oxygen species (ROS) are also affected by βHB; it alters mitochondrial metabolism via the NAD+/NADH and ubiquinone/ubiquinol ratios, therefore decreasing ROS.

Molecular patterns associated with pathogens and damage (PAMPs and DAMPs) enhance the production of NLRP3, an inflammasome that leads to the activation of proinflammatory interleukins such as IL-1β and IL-18, and is inhibited by βHB (36). Finally, βHB increases the acetylation of histones, leading to a higher expression of oxidative stress suppression genes, activation of anti-inflammatory interleukins like IL-4, and conversion of macrophages from M1 (proinflammatory) to M2 (anti-inflammatory), Ganggand et al. also reported low levels of IL-1β and TNF-α in the hippocampus of mice under stress after preadministration of βHB.

Our results also showed that NLRP3 was lower when 1-3 Hydroxybutanediol Monoester was administered, when endogenous ketogenesis is stimulated. Therefore, there exists an inhibition of pro-inflammatory cytokines; such as TNF-1, IL-1β, related to the activation of NLRP3, and this is directly related to the down-regulation of NFkB in endoK.

In the context of SCI, Nrf2 plays a key role in multiple cellular processes, such as regulating apoptosis, modulating inflammatory responses, suppressing the NF-κB pathway, maintaining iron homeostasis, promoting autophagy, supporting axonal regeneration, and preserving mitochondrial function. According to Mao et al., mice lacking Nrf2 after SCI exhibited greater motor impairments, increased neuronal death, delayed responses in electrophysiological assessments, elevated expression of pro-inflammatory cytokines IL-6 and IL-1, and reduced levels of antioxidant and detoxification enzymes, including NQO1 and GST-1 (42) The reason for this is that oxidative stress is a key factor for the perpetuation of the secondary injury, and Nrf2 promotes the transcription of antioxidant response elements (42), regulating the expression of up to approximately 500 genes. These include redox-regulating factors such as heme oxygenase-1 (HO-1), superoxide dismutase 1 (SOD1), and glutathione peroxidase 1 (GPx1), as well as detoxification enzymes and various stress-response proteins (*44*)

As reported by Yao Lu, in rats who suffered SCI and were fed with a ketone diet, NRf2 levels were statistically significantly increased compared to the group that was not fed with a ketone diet (44) This, as addressed before, leads to an attenuation of inflammation via decreasing NFkB pathways and expression of chemokines such as IL-1β, TNF-α, IFNγ, and myeloperoxidase activity. Acting as a synergistic anti-inflammatory pathway alongside the decrease of NLRP3 after βOHB also inhibits NFkB (46)

In this study, the administration of 1-3 Hydroxybutanediol Monoester and the use of a KD conferred partial protection against SCI secondary injury, clearly affected pro-inflammatory chemokine concentration.

## Conclusion

In summary, our results show that either endogenous or exogenous ketosis (1-3 Hydroxybutanediol) provoked a positive effect in spinal cord injury induced in Wistar rats in the first eight hours of that injury. Ketone bodies act as an antioxidant agent, due to the increase in SOD2, catalase through histone acetylation, G6PDH, NQO1, γGCLM, and SOD2 by Nrf2; therefore, the inflammation response was lower. Furthermore, the inflammasome NLRP-3 decreased, leading to a decrease in the levels of TNF-α and IL-β, which are closely related to the expression of NLRP-3,. When spinal cord injury exists, not only is an inflammatory response activated, but also an ischemia phenomenon appears, and HIF-1 is overexpressed. This study shows that the ketone bodies act by stimulating the succinate molecule, leading to an increase in HIF-1α and then the over-transcription of NFkB. Nevertheless, some proinflammatory cytokines are still at lower levels, for instance, IL-1β and TNF-α. It is worth noting that endoK was stimulated for twenty-four hours before the spinal cord injury was carried out, and exogenous ketosis was induced after the spinal cord injury took place (two and five hours).

Finally, we must keep in mind that we are studying a model of acute spinal cord injury and more research is needed to elucidate if the dosage of 1-3 Hydroxybutanediol could be modified, either by adjusting the time administration or dosage. Altough endogenous ketosis showed better results, exogenous ketosis could help with the control of secondary injury because we can manage its use after SCI.

## Acknowledgements.

I thank technicians Miguel García and Victor Gómez for their support during the surgical procedures performed under this protocol.

## Data Availability

All relevant data are within the manuscript and its Supporting Information files.

## Author contributions

Conceptualization: G.R.S., C.C.B.R., A.S.M.

Methodology: A.C., P.M.R., J.A.M., M.V., C.B.

Investigation: A.C., P.M.R., J.A.M., M.V.

Laboratory analysis: A.C.

Histopathology development: C.B.

Histopathology and immunohistochemistry evaluation: R.V.

Data curation: M.A.P.C., M.G.P.S.

Formal analysis: A.S.M., M.A.P.C., M.G.P.S.

Writing – original draft: A.C.

Writing – review & editing: A.S.M., V.R., G.R.S., C.C.B.R.

Supervision: A.S.M., V.R., G.R.S., C.C.B.R.

